# A Hyperbolic Discrete Diffusion 3D RNA Inverse Folding Model for functional RNA design

**DOI:** 10.1101/2025.03.11.642534

**Authors:** Dongyue Hou, Shuai Zhang, Mengyao Ma, Hanbo Lin, Zheng Wan, Hui Zhao, Ruian Zhou, Xiao He, Xian Wei, Dianwen Ju, Xian Zeng

**Affiliations:** Department of Biological Medicines and Shanghai Engineering Research Center of Immunotherapeutics, School of Pharmacy, Fudan University, Shanghai 201203, China; Byterna therapeutics, Shanghai 201203, China; MoE Engineering Research Center of Hardware/Software Co-design Technology and Application, East China Normal University, Shanghai 200062, China; Shanghai Engineering Research Center of Molecular Therapeutics and New Drug Development, Shanghai Frontiers Science Center of Molecule Intelligent Syntheses, School of Chemistry and Molecular Engineering, East China Normal University, Shanghai 200062, China; Faculty of Sciences, Engineering and Technology, the University of Adelaide, Adelaide, South Australia, 5070, Australia; Shanghai Engineering Research Center of Molecular Therapeutics and New Drug Development, Shanghai Frontiers Science Center of Molecule Intelligent Syntheses, School of Chemistry and Molecular Engineering, East China Normal University, 500 Dongchuan Road, Shanghai, 200062, China; Chongqing Key Laboratory of Precision Optics, Chongqing Institute of East China Normal University, Chongqing, 401120, China; New York University-East China Normal University Center for Computational Chemistry, School of Chemistry and Molecular Engineering, New York University Shanghai, Shanghai, 200062, China

**Keywords:** RNA inverse folding, generative RNA design, hyperbolic embedding, discrete diffusion model

## Abstract

Generative design of functional RNAs presents revolutionary opportunities for diverse RNA-based biotechnologies and biomedical applications. To this end, RNA inverse folding is a promising strategy for generatively designing new RNA sequences that can fold into desired topological structures. However, three-dimensional (3D) RNA inverse folding remains highly challenging due to limited availability of experimentally derived 3D structural data and unique characteristics of RNA 3D structures. In this study, we propose RIdiffusion, a hyperbolic denoising diffusion generative RNA inverse folding model, for 3D RNA design tasks. By embedding geometric features of RNA 3D structures and topological properties into hyperbolic space, RIdiffusion efficiently recovers the distribution of nucleotides for targeted RNA 3D structures based on limited training samples using a discrete diffusion model. We perform extensive evaluations on RIdiffusion using different datasets and strict data-splitting strategies and the results demonstrate that RIdiffusion consistently outperforms baseline generative models for RNA inverse folding. This study introduces RIdiffusion as a powerful tool for the generative design of functional RNAs, even in structure-data-scarce scenarios. By leveraging geometric deep learning, RIdiffusion enhances performance and holds promise for diverse downstream applications.

## INTRODUCTION

Engineered functional ribonucleic acids (RNAs) are now widely applied across various biomedical fields, including RNA-based gene editing, vaccines, therapeutics, and synthetic biology elements.^1,2^ Properly folded RNA molecules exhibit a broad range of biological functions by interacting with other molecules (e.g., small compounds, DNA, and proteins), and these functions are often driven by their 3D structures.^3–5^ Therefore, designing nucleotide sequences that can fold into specific 3D structures holds the promise of creating novel functional RNAs, potentially yielding profound impacts on RNA biomedical applications.^6,7^ In recent years, generative deep learning algorithms have made a series of breakthroughs in the modelling and design of protein and small molecules.^8^ Although advancements in RNA applications are still in their infancy, structure-based generative design algorithms offer unique opportunities for functional RNA design, potentially unlocking the full potential of RNA technologies across various biomedical applications.^9–11^

The primary goal of RNA inverse folding is to design nucleotide sequences that can fold into a given RNA secondary or tertiary structure. However, existing tools either focus on 2D inverse folding tasks or learn limited 3D structural features from experimentally determined and predicted 3D structure datasets for the 3D inverse folding problem.^12–15^ Recently, several studies^12,16,17^ have attempted to address the RNA 3D inverse folding task.

The most serious bottleneck behind RNA 3D-related modeling tasks is the scarcity of experimentally derived RNA 3D structural data. High-resolution RNA structures are significantly less common compared to those of proteins.^18^ In addition, the potential benefits of synthetic 3D structural data for 3D structure-related AI modelling is undefined, given the fact that current models struggle to achieve high accuracy in RNA 3D structure prediction.^19–22^ The “low-sample” challenge underscores the importance of learning efficiency in generative models in 3D RNA inverse folding tasks. Therefore, it is essential to develop more efficient algorithms to address the unique challenges of the 3D RNA inverse folding problem.

Diffusion models have largely advanced the high-quality generation of proteins and small molecules.^23–25^ In this study, we interpret the inverse folding process as a denoising problem in the diffusion model, and employ discrete noise space to iteratively predict nucleotide base type (A, U, G or C) from RNA structure backbone for RNA sequence generation. In addition, RNA structures exhibit inherent hierarchical organization (e.g., from base-paring, secondary structure elements and remote pseudoknots to complex 3D shapes),^26^ which may be more effectively represented in a hyperbolic space. Recently, we and others have revealed that, compared to Euclidean space, a hyperbolic distance better captures molecular differences and improve performance in small molecule generation tasks.^27,28^ Hyperbolic space may be advantageous for representing subtle structural variations in RNAs, thereby enhancing the representation learning of RNA 3D structural information to alleviate the “low-sample” challenge (Figure 1A, 1B, 1D). Therefore, in this study, we propose RIdiffusion (RNA Inverse-folding diffusion), which employs hyperbolic equivariant graph neural networks (HEGNNs) to parameterize the discrete diffusion model, and effectively capture the 3D structural characteristics by incorporating RNA’s geometric features and topological properties into the generation process.

**Figure 1.**
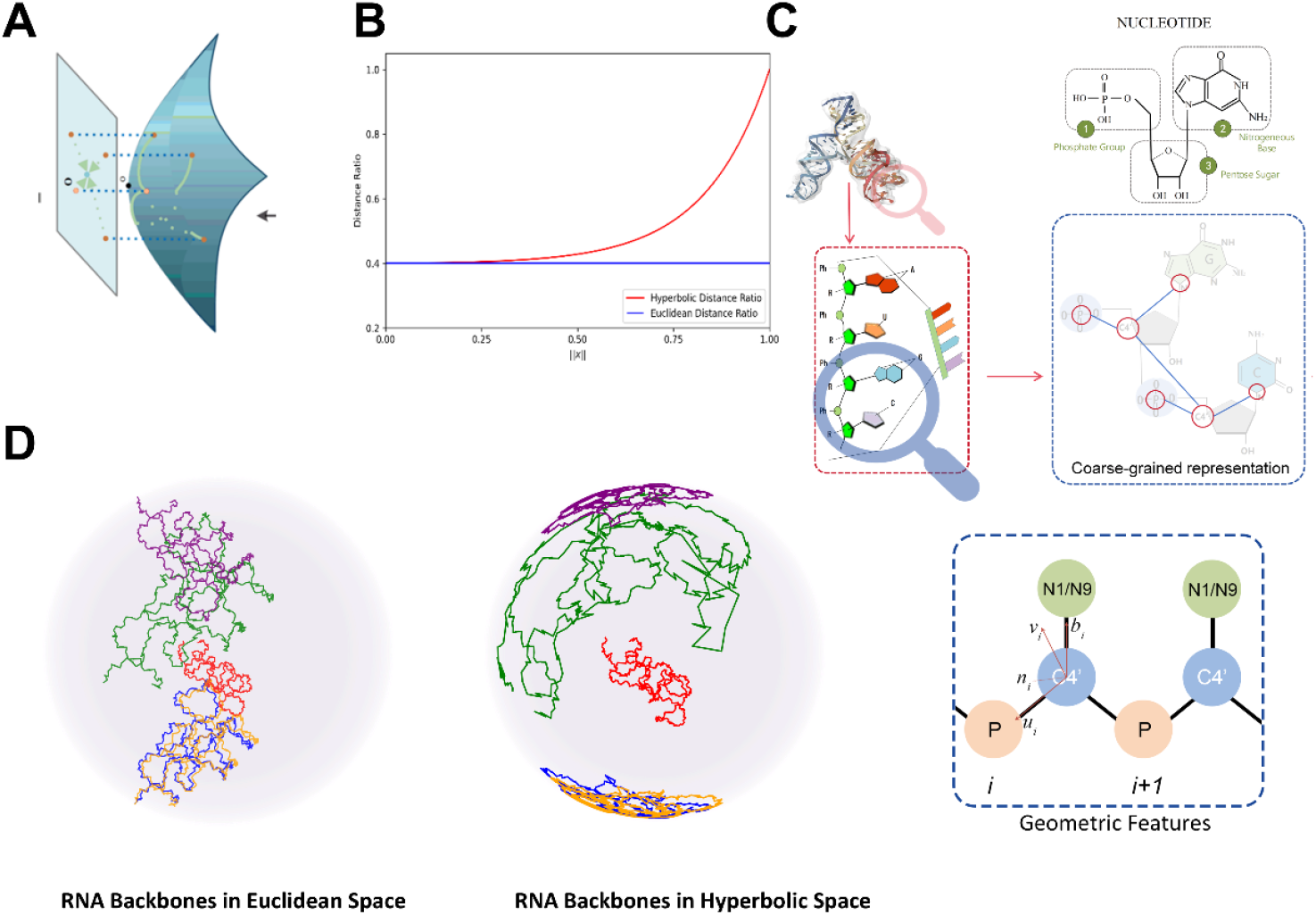
Hyperbolic space represents RNA 3D backbones in a more efficient pattern compared to Euclidean space. (A) Illustration of embedding graph to hyperbolic space. (B) In hyperbolic space, the distance ratio 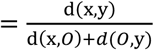 between vertices (where both x and y vertices are equidistant from origin *O*) increases, making it a more accurate representation of graph-like structures, compared to Euclidean space. ||x|| denotes the distance from a given vertex x to *O*. (C) Chemical components, 3D structures and geometric features of RNA. C4’, P and N1/N9 atoms are utilized for coarse-grained representation and geometric features calculation (see details in Figure S1, Table S1). (D) Mapping the RNA backbones from Euclidean space to the Hyperbolic space (Poincaré Ball) better captures and distinguishes structural variations.

To comprehensively evaluate RIdiffusion, we constructed multiple datasets by partitioning the Protein Data Bank (PDB)^29^ and RNAsolo dataset^30^ based on sequence length and similarity thresholds. Results demonstrated that, on low-sequence-similarity datasets, RIdiffusion outperformed the second-best model by 7.27%, and consistently surpassed both benchmark and state-of-the-art (SOTA) models across multiple benchmarks. Our ablation studies also indicated that hyperbolic embedding and equivariant graph transformer significantly enhanced the performance of 3D RNA inverse folding. In summary, this study establishes RIdiffusion as a powerful tool for designing novel sequences that fold into desired 3D RNA structures (Figure 2A), unlocking new opportunities for functional RNA design.

**Figure 2.**
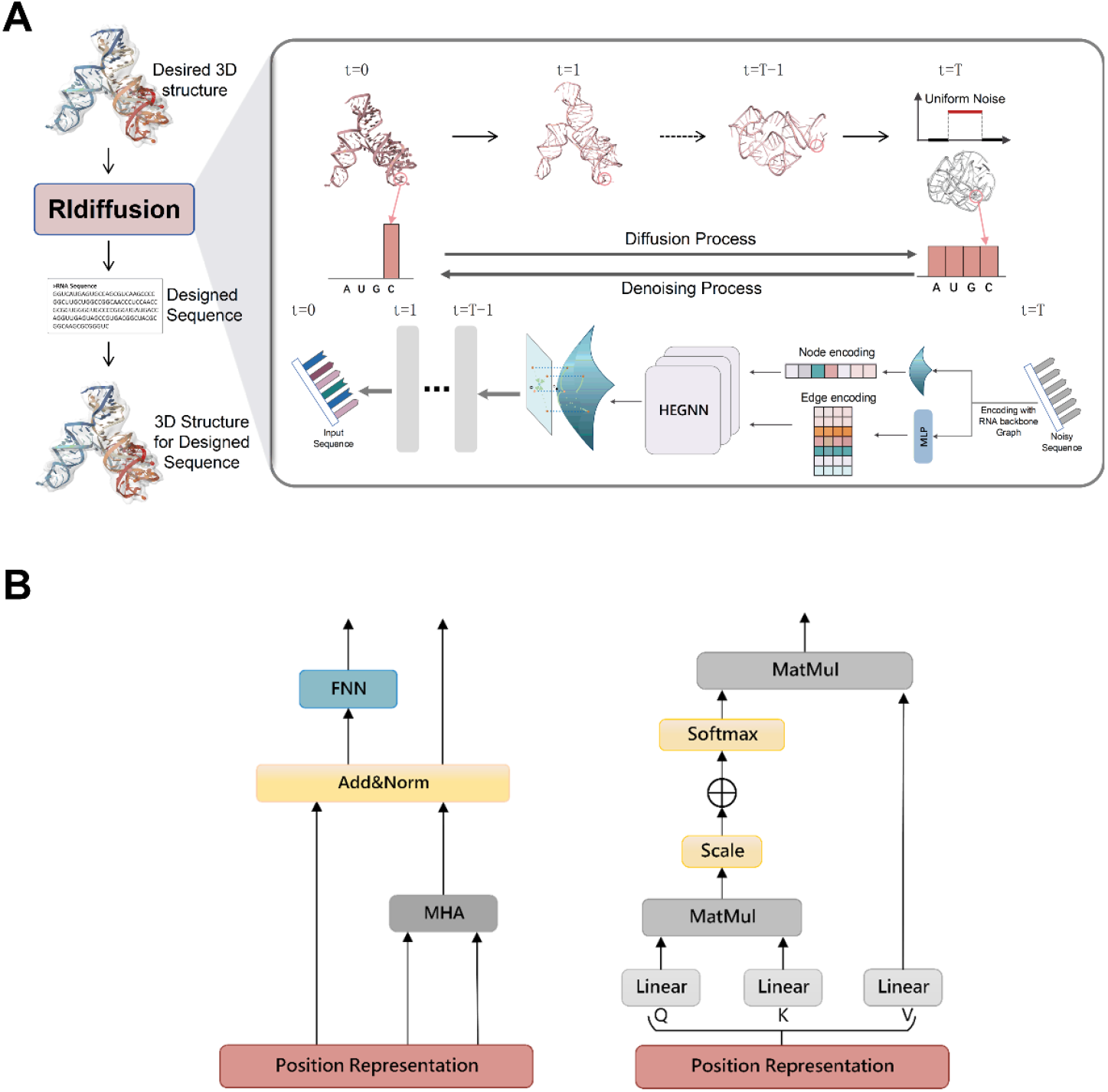
**(A)** Overview of RIdiffusion. During the diffusion process, uniformly distributed noise is progressively added to the nucleotide sequence of a real RNA molecule, causing the structure represented by the sequence to undergo random transformations until the sequence distribution completely transitions into uniformly distributed noise. In the denoising process, the nucleotide sequence is first randomly sampled from a uniform distribution and then gradually denoised through a neural network, ultimately generating a realistic nucleotide sequence. **(B)** Architecture of the modified Transformer module for encoding residue positional information in hyperbolic space. The figure illustrates the design of the modified Transformer block used to encode the positional information of RNA residues and capture their global relationships (left panel). In this architecture, the initial residue positional information is processed through several steps: Multi-head Attention (MHA), layer normalization, Feedforward Neural Network (FFN). The figure also shows the single-head attention function (right panel), detailing the attention mechanism that computes residue relationships.

## MATERIALS AND METHODS

### Residue Graph of Folded RNA

The geometric structure of RNA is crucial for its function. To capture the complex 3D geometric information of folded RNA molecules, we represent the 3D structure of RNA using a k-nearest neighbor graph G = (**X, E**), where the nodes correspond to C4’, P, and N atoms (N1 for pyrimidines or N9 for purines) (Figure 1C). This coarse-grained backbone representation encodes both the physicochemical properties and geometric details of the RNA. Node features **X** = [X^nt^, X^pc^, X^di^, X^mur^, X^pos^] include nucleotide types **X**^nt^, physicochemical properties **X**^pc^, and local environment information [**X**^di^, **X**^mur^]. The physicochemical properties **X**^pc^ consist of the Solvent Accessible Surface Area (SASA), which measures the extent to which nucleotides are exposed to solvent in the RNA molecule, and the B-factor. Local environment information includes RNA surface-aware^31^ feature **X**^mur^, which are nonlinear projections of the weighted average distance from the central nucleotide to its one-hop neighbors. Additionally, the dihedral angles **X**^di^ of backbone atoms, which describe the conformation of the RNA backbone and are represented using sine and cosine functions and **X**^pos^ denotes the 3D coordinates of residue. We perform diffusion on the nucleotide features **X**^nt^ and denoise them using HEGNN.

### Edge features

**E** = [E^rbf^, E^lf^, E^cont^, E^sr^] contains the relationships between connected nodes, determined by encoding schemes such as relative spatial distances and sequential distances. For adjacent residues *i* and *j*, their kernel-based distance **E**^rbf^ is determined by a Gaussian Radial Basis Function (RBF). The relative positions of residues **E**^lf^ are defined by local frames^31^ created from backbone atoms. We determine whether two residues are in contact^32^ **E**^cont^ in space based on the cutoff distance between nodes and residues’ sequential relationship **E**^sr^ encoded by their relative position.

### The Poincaré Ball Model

Riemannian geometry^33,34^ is a powerful mathematical branch that studies the properties of manifolds equipped with a Riemannian metric. It is widely used to address non-Euclidean geometric problems in machine learning. A Riemannian manifold^35^ is a smooth manifold equipped with a Riemannian metric, denoted as (ℳ, g). At each point x ∈ ℳ, there is a tangent space 𝒯_*x*_ℳ. The Riemannian metric g on ℳ is a collection of inner-product functions g_x_: T_x_M × 𝒯_x_ℳ → ℝ defined on these tangent spaces. The Riemannian metric is used to measure distances on the manifold.^36^

Hyperbolic space^37^ refers to a Riemannian manifold with constant negative curvature. The Poincaré disk ball 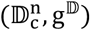 is one of the fundamental mathematical models^38–43^ representing hyperbolic geometry. It characterizes the properties of hyperbolic geometry by mapping the entire hyperbolic plane into the interior of a disk, defined by the manifold 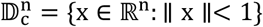 with the Riemannian metric: 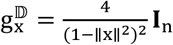 and Poincaré Ball of radius c.

Distance: For the given points x, y ∈ 𝔻^n^,the induced distance can be written by:

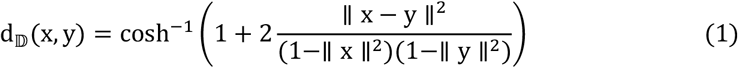

In hyperbolic space, d_𝔻_(x, y) more closely resembles the graph distance as x and y approach the edge of the Poincaré ball, making it a better representation of graph-like structures compared to Euclidean space. For example, consider two vertices x and y as children of a parent z in a tree, with z placed at the origin. The graph distance between x and y is the sum of the distances from x to z and from y to z, i.e., d(x, y) = d(x, *O*) + *d*(*O*, y). Normalizing this gives the distance 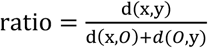. In Euclidean space, the distance ratio remains constant, which means it fails to capture the graph’s structure effectively. In contrast, in hyperbolic space, as the distance increases, the hyperbolic distance approaches the graph distance, with the distance ratio rising (Figure 1B). This makes hyperbolic space more suitable for representing tree-like structures.

Exponential and logarithmic maps: In hyperbolic space, the exponential map and the logarithmic map are mathematical tools used to convert points between the hyperbolic space and its tangent space. These maps facilitate geometric operations between hyperbolic space and Euclidean space.

For 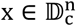, the exponential map expc: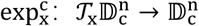 assigns to a tangent vector 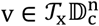 the point 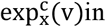 hyperbolic space is given by:

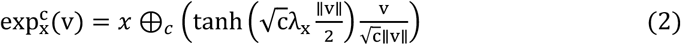

The logarithmic map is the inverse operation of the exponential map. It maps a point *x* in hyperbolic space back to a vector *v* in the tangent space. The logarithm map is given by:

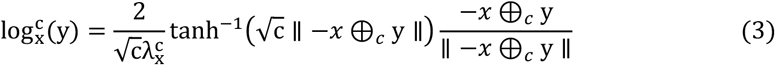

where ⊕_*c*_ is Möbius addition. ^44^

Parallel transport: In hyperbolic geometry, parallel transport is an operation that moves a vector from one point to another while preserving the vector’s angle and length relative to its starting point. Specifically, parallel transport is a transformation that maintains the Riemannian metric tensor, which includes norms and inner products. Given a smooth manifold ℳ, parallel transport P_x→y_(·) maps a vector **v** ∈ 𝒯_x_ℳ to P_x→y_(**v**) ∈ 𝒯_y_ℳ.

### Feature Representation in Hyperbolic Space

Hyperbolic space possesses higher representational capacity and efficient distance computation properties, making it particularly useful for molecular embeddings. Firstly, hyperbolic space can capture complex structures in high-dimensional space at lower dimensions, thus reducing information loss during the embedding process. This is crucial for molecules with diverse and complex structures. Secondly, the distance computation characteristics in hyperbolic space allow it to represent better and calculate similarities between molecules. We adapt a hyperbolic encoder^45,46^ to generate hyperbolic embedding.

We first map Euclidean node features x^E^ to the Poincaré Ball 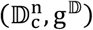 via the exponential map. Let 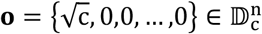 denote the origin in 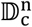 (Figure 1D).

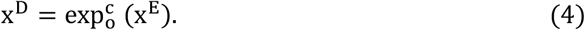

Subsequently, we perform a linear transformation after projecting the hyperbolic point ***X***^***D***^ to the tangent space 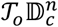 using the logarithmic map and use the exponential map to project the result back to the manifold. Finally, we use σ to keep manifold preserving.

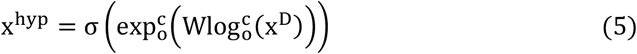

### Hyperbolic Positional Encoding

In the previous context, we utilized various encodings for the local environment of residue positional information. While local environmental information is vital for learning residue representations, global environmental information also plays a significant role. To capture the impact of global information on residues, we employ a modified Transformer module to encode the positional information **X**^pos^ of residues to capture the global relationships between them and map them to hyperbolic space. The modified Transformer^47^ block, as illustrated in Figure 2B (left), is defined as follows:

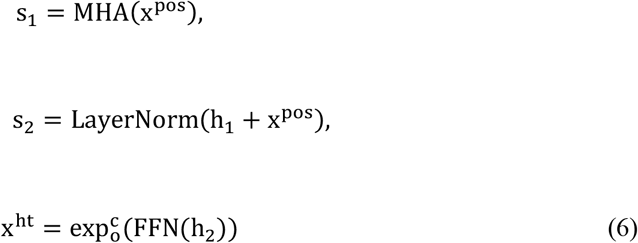

where MHA denotes the multi-head attention layer (the definition of the single-head attention function is shown on the right of Figure B), and FFN is a feedforward neural network

### Discrete Diffusion Model

The Discrete Diffusion Model learns the complex distribution of data by progressively adding noise to simulated data and performing reverse denoising, thereby generating high-quality new samples.

#### Diffusion process

In the forward process, noise is continuously added to the data until it is transformed into pure noise. This process can be represented by a transition matrix, which facilitates the transition from the current time step to the next. The transition matrix is iteratively applied to the original data, which evolves over time due to the influence of noise. As the number of time steps increases, the original data eventually converges towards a uniform distribution across all data types. For discrete variables with n types, the transition probabilities can be defined by the matrix [**Q**_*t*_]_*ij*_ = *q*(*x*_*t*_ = *j* ∣ *x*_*t* − 1_ = *i*). Based on **Q**_*t*_ we can set the following transition kernel:

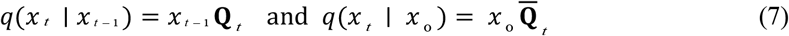

where 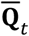 is an accumulating version of **Q**_*t*_.

#### Parameterization of the denoising process

The reverse process aims to restore the original data distribution by progressively removing noise. This process iteratively refines the noisy data back to its original form. The reverse process is guided by learned denoising functions, which predict the denoised data at each step, aiming to approximate the inverse of the forward process. New samples can be iteratively generated through reverse diffusion. The probability distribution p_θ_(x_t−1_ ∣ x_t_) is obtained from the predicted probability 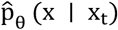 by the neural network. We obtain the following parameterization through the marginalization of the network predictions:

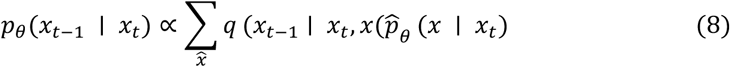

The posterior^48^ can be written as:

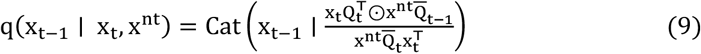

where Cat is a categorical distribution of x.

### Hyperbolic Equivariant Graph Denoising Network

Biomolecules such as RNA have complex 3D structures, which are crucial for their function. Any rotation or translation of the RNA model in three-dimensional space should not alter its intrinsic properties and functions. Traditional neural networks may lose important geometric information when processing 3D data, leading to inaccurate or unreliable predictions of RNA function. To address this issue, we utilize SE(3) equivariant neural layers from Equivariant Graph Neural Networks (EGNN).^49^ This approach allows the model to preserve SO(3) rotational equivariance and E(3) translational invariance when updating the representations of nodes and edges, thereby retaining geometric information and ensuring the robustness and effectiveness of the hidden representations, to enable the EGNN network to update edge.

For representations and operations in hyperbolic space, we propose HEGNN, which is composed of Hyperbolic Equivariant Graph Convolutional Layers (HEGCL). HEGCL is defined as follows to update the hidden representations of nodes:

To update the new edge embeddings 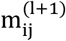,we first use the logarithmic map 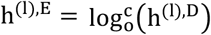 to project the hyperbolic representations of the node hidden Representationh 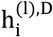 (with initial representation x^hyp^) into the tangent space 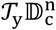.Then, using the hyperbolic encoding distanced 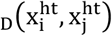 from Equation 1 and the current edge embedding 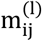,the new edge embeddings can be defined as follows:

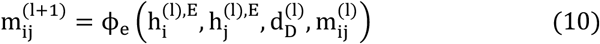

The new hyperbolic position encoding 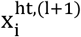 can be updated based on the current Euclidean node encoding 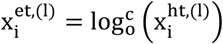:

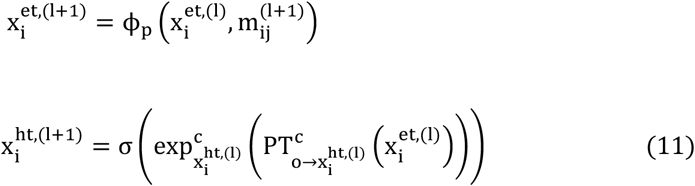

Based on the above, the update of the new node features 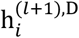 can be represented in the following form:

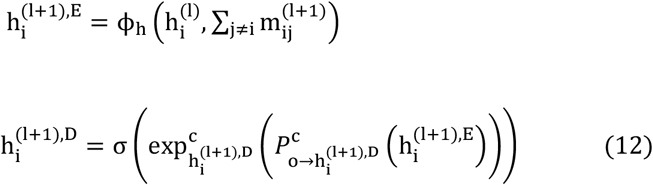

where *ϕ*_*h*_, *ϕ*_*p*_, *ϕ*_*e*_ are learnable function parametrized by neural networks.

### Training Denoising Networks and DDIM Sampling

We aim to simulate the nucleotide sequence diversity of RNA while preserving its inherent structural constraints. We train the denoising neural network for optimizing the predicted probability 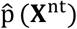 of each node’s nucleotide type, the training process is shown in Algorithm 1. The algorithm takes an RNA graph with node features and edge features, including physicochemical properties and geometric attributes, as input. A time step tis sampled from a uniform distribution, and then the cumulative transition probability is calculated to generate noisy data and embed it into hyperbolic space. The noisy data is predicted by the denoising network HEGNN.

#### Algorithm 1 Training

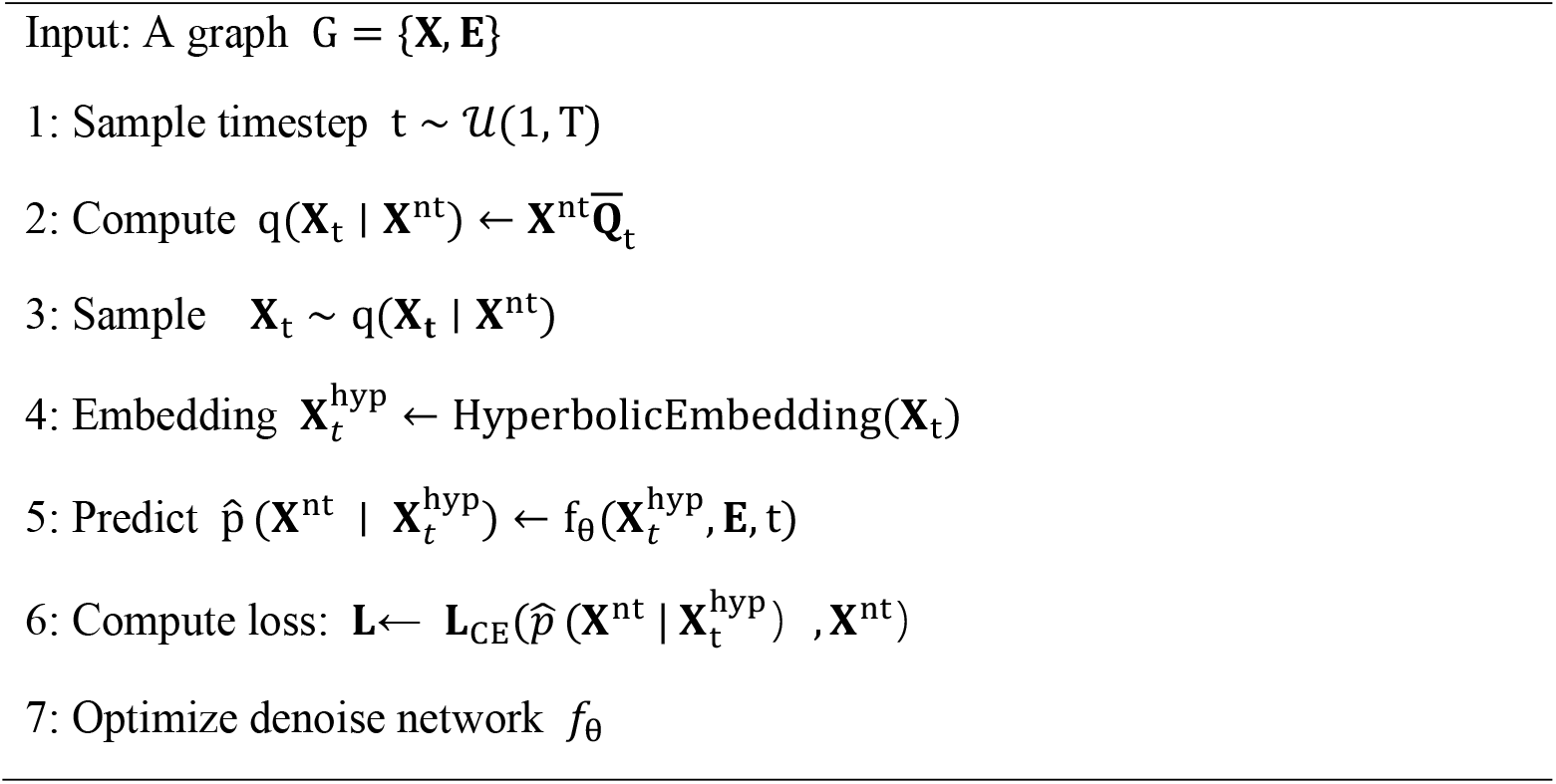

Since we possess the closed-form expressions for the predicted **X**^nt^ and the posterior distribution p(x_t−1_ ∣ x_t_, x^nt^)described in Equation 9. After completing the training, we can sample data using the neural network and the posterior distribution. DDIM^50^ significantly speeds up the data generation process by using a non-Markovian forward diffusion process, reducing the number of incremental steps. By setting the noise variance to zero at each step, DDIM makes the reverse generation process fully deterministic and repeatable. We describe the DDIM sampling process in Algorithm 2.

#### Algorithm 2 DDIM Sampling

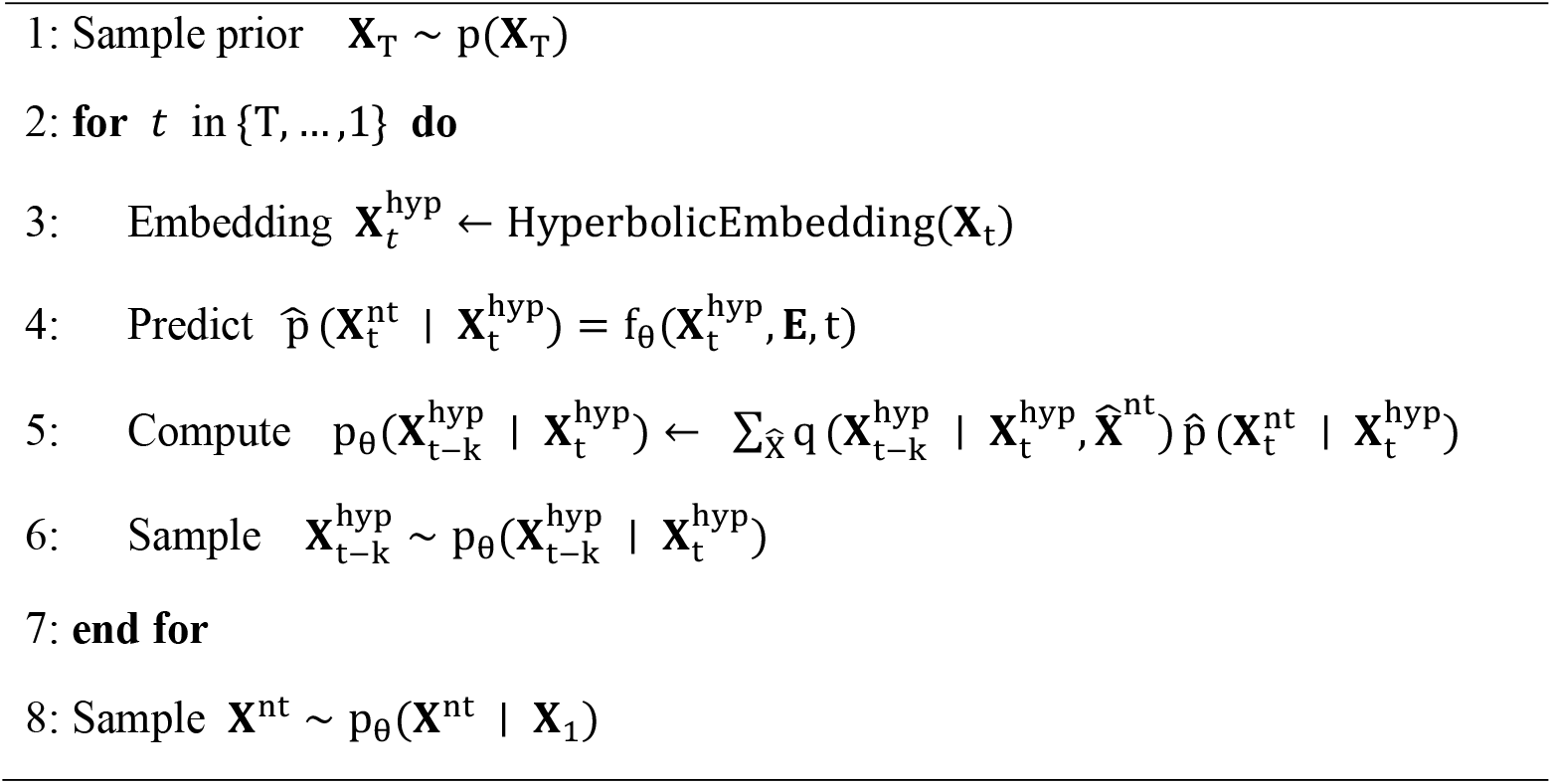

### Predicting and embedding RNA Secondary Structure (SS)

To enrich the RNA secondary structure (SS) information, we predicted base-paring information using RNAstructure,^51^ EternaFold,^52^ and MultiRNAFold.^53^ Next, we employed bpRNA^54^ to consistently annotate eight SS types: bulges, hairpin loops, multiloops, internal loops, pseudoknots, exterior loops, stems, and segments. Finally, the SS information was converted into one-hot embeddings and processed by RIdiffusion.

### Datasets and Setup

We trained our model using the publicly available PDB^29^ and RNAsolo dataset.^30^ The RNAsolo dataset, sourced from public databases, includes a large collection of RNA sequences and experimentally determined RNA structures. These structures are categorized into Solo RNA, protein-RNA complexes, and DNA-RNA complexes, each with varying levels of resolution (Table S2 and S3). We followed the same splitting methods outlined in RDesign^14^ to ensure a high-quality dataset with only experimentally derived structures, which consists of 1773 structures for training, 221 structures for validation, and 223 structures for testing in an 8:1:1 ratio. As sequence similarity increases, the likelihood of structural similarity also grows. To evaluate the model’s generalization capability, we further used PSI-CD-HIT^55^ to cluster the sequences based on nucleotide similarity, setting similarity thresholds of 0.8, 0.9, and 1.0. This process generated three subsets: seq-0.8, seq-0.9, and seq-1.0.

### Evaluation Metrics

We use recovery rate (RR) and Macro-F1 score as indicators to evaluate the accuracy of the generated nucleotide sequences. The RR is the most commonly used metric in inverse folding, which calculates the positional consistency between the model-predicted sequence and the actual sequence. It has a higher likelihood of achieving correct folding for similar sequences. The Macro-F1 score is used to assess the accuracy and comprehensiveness of the model in predicting protein or RNA sequences, particularly in cases where the distribution of different letter residues is imbalanced, by evaluating the Macro-F1 score for each residue in the sequence.

The Novelty Score is employed to quantify the degree of difference and uniqueness between the generated samples and those in the training dataset. It is defined as follows:

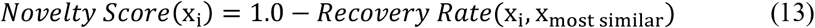

where: x_i_ represents the i-th generated sequence; x_most similar_ denotes the RNA sequence in the training set that exhibits the highest RR when compared to x_i_.

To assess the overall novelty relative to the training space for each model, we defined the global-scale novelty (GSN) score as follows: First, we generated batches of samples using different algorithms, with each original structure in the given test dataset producing 100 sample sequences. For each batch of structure-guided samples, we selected sequences with maximum, medium, and minimum RRs and calculated their novelty scores relative to the training dataset separately. Finally, we categorized the sequences into maximum, medium, and minimum groups, and computed the mean novelty score for each group, thereby deriving the GSN score across these three levels. High GSN score indicates that the generated samples are broadly distributed in the feature space, exhibiting high diversity and innovation. Meanwhile, RNAincoder^56^ was utilized to generate embeddings for t-SNE visualization in physicochemical novelty evaluation.

In the foldability evaluation section, for each case study, we generated 10 sample sequences based on the original given structures. We then performed *in silico* RNA tertiary structure prediction using RhoFold.^19^ Additionally, 3D structure alignment software US-align^57^ was used to align the predicted structures and calculate their RMSD.

### Baselines

To evaluate the effectiveness of our proposed method, we compare it with one RNA tertiary structure inverse folding model, three protein tertiary structure inverse folding baselines, and one Transformer-based graph GNN network for graph structure: gRNAde,^16^ PiFold,^58^ StructGNN,^32^ GVP-GNN^59^, and GraphTrans.^60^ Since the training scripts of RDesign^14^, RiboDiffusion^12^, and RhoDesign^17^ were not available, conducting a fair comparison was challenging. Therefore, we compared our model with open-source inverse folding models, using their default hyperparameters to ensure fairness. For this comparison, we trained RIdiffusion and re-trained baseline models on the same dataset (as previously described) while adhering to the default hyperparameters of the respective baseline models.

## RESULTS

### The architecture of RIdiffusion

Compared to traditional diffusion models,^50,61,62^ discrete diffusion models^48^ are particularly well-suited for handling discrete data, such as textual data like nucleotide sequences. Mechanistically, discrete diffusion more effectively captures the structural information of discrete data by defining more structured discrete corruption processes, such as simulating Gaussian kernels and embedding spaces based on nearest neighbors.^48^ Therefore, we adopted discrete diffusion model^48^ and one of its variants^63^ in RIdiffusion (Figure 2A). The quality of the generated sequences in diffusion models is largely determined by the denoising network. In this section, we propose a hyperbolic-based denoising network, which embeds molecular structural features into an equivariant hyperbolic space to capture molecular hierarchical structures and structural variations more sensitively (Figure 2B). Notably, different RNA backbones are separated more obviously in a hyperbolic space than in the Euclidean space (Figure 1D).

### RNA 3D inverse folding performance of RIdiffusion

Several generative models have recently been proposed for inverse folding tasks. For example, RDesign,^14^ gRNAde,^16^ and RiboDiffusion^12^ were proposed for RNA 3D inverse folding, while GVP-GNN,^59^ PiFold,^58^ and StructGNN^32^ are representative tools for protein 3D inverse folding. Additionally, GraphTrans^60^ was also included as a Transformer-based graph GNN network for graph structure. To ensure fair comparisons, we included all these models except for RDesign, RiboDiffusion, and RhoDesign due to the unavailability of its training codes. As shown in Table 1, RIdiffusion achieved the highest RRs and Macro-F1 scores in all similarity-cutoff datasets, when compared with baselines. Interestingly, RIdiffusion outperformed gRNAde in Macro-F1 scores by 11.76%, 16.34% and 17.46% at seq-0.8, seq-0.9 and seq-1.0 datasets, respectively. Among those baselines, PiFold and StructGNN consistently outperformed the remaining ones. Surprisingly, RIdiffusion, trained on the same dataset (seq-1.0) as RDesign, slightly surpassed RDesign in both RR and Macro-F1 score, with RDesign’s reported mean values of 41.53% and 0.4089, respectively.

**Table 1.**
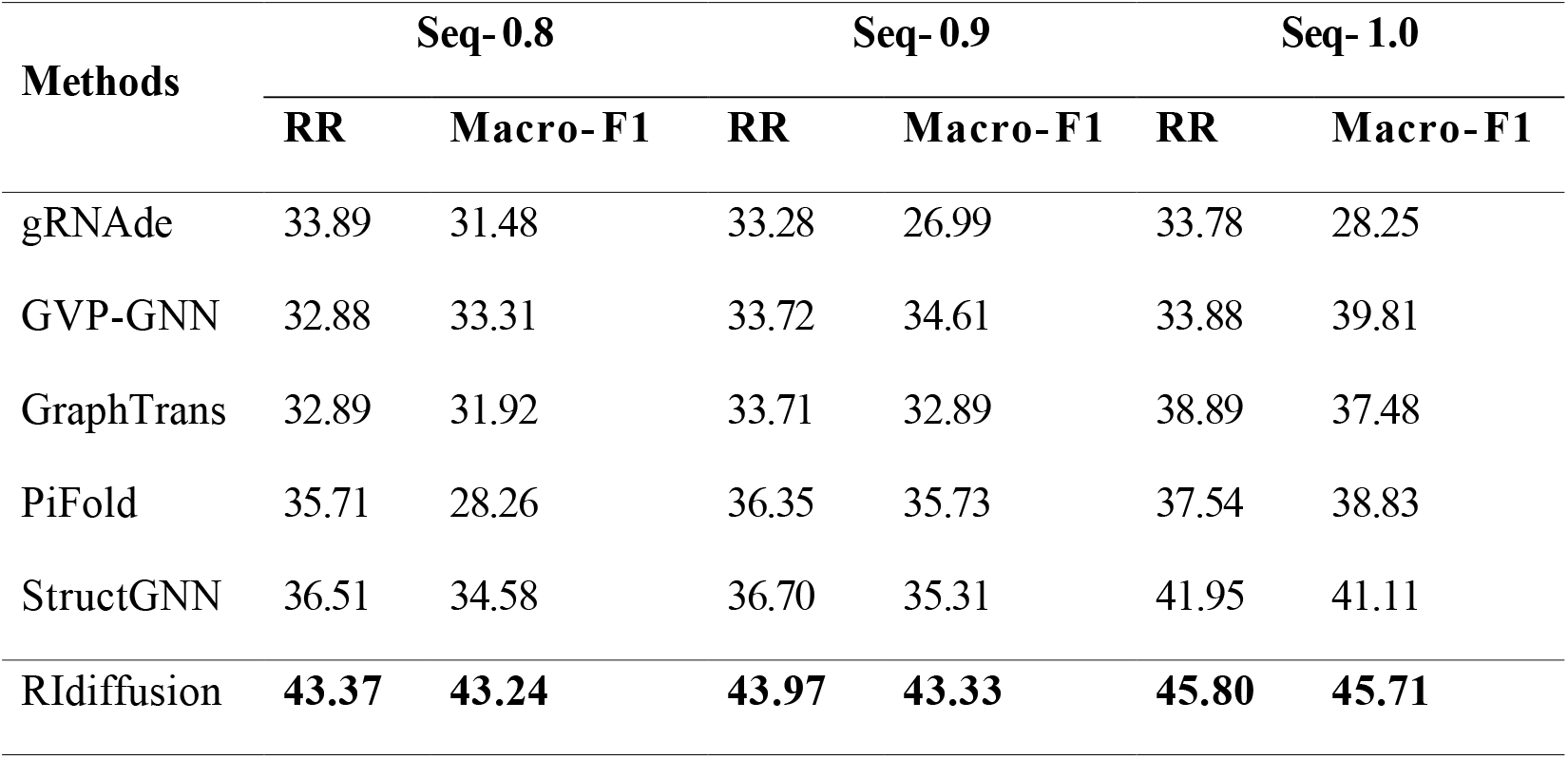
Comparison of inverse folding performance across methods, showing the sequence recovery rate (RR, %) and Macro-F1 score (Macro-F1, × 100) at different sequence similarity thresholds (seq-0.8/0.9/1.0).

To further analyze the impact of RNA sequence length on performance, we divided the dataset of seq-0.8 into short (<50nt), medium (50∼100nt), and long (>100nt) sequence subsets to train and test RIdiffusion again. The results showed that RIdiffusion consistently outperformed the baselines in all 3 subsets (Table 2). The medium subset resulted in a superior RR and Macro-F1 scores, compared to short and long sequence subsets, which is consistent with the findings of RiboDiffusion^12^. Interestingly, the medium subset of seq-0.8 dataset achieved higher performance than the original seq-0.8 dataset, suggesting that the distribution of sequence lengths may impact generation quality. These results also suggest that RIdiffusion seems not sensitive to the decrease in sample size, given the fact that seq-0.8-medium achieves RR of 45.43% on 225 training samples while seq-0.8 achieves RR of 43.37% on 761 training samples. Collectively, our data suggest that RIdiffusion exhibits superior performance on RNA 3D inverse folding task, even trained on limited structural samples.

**Table 2.**
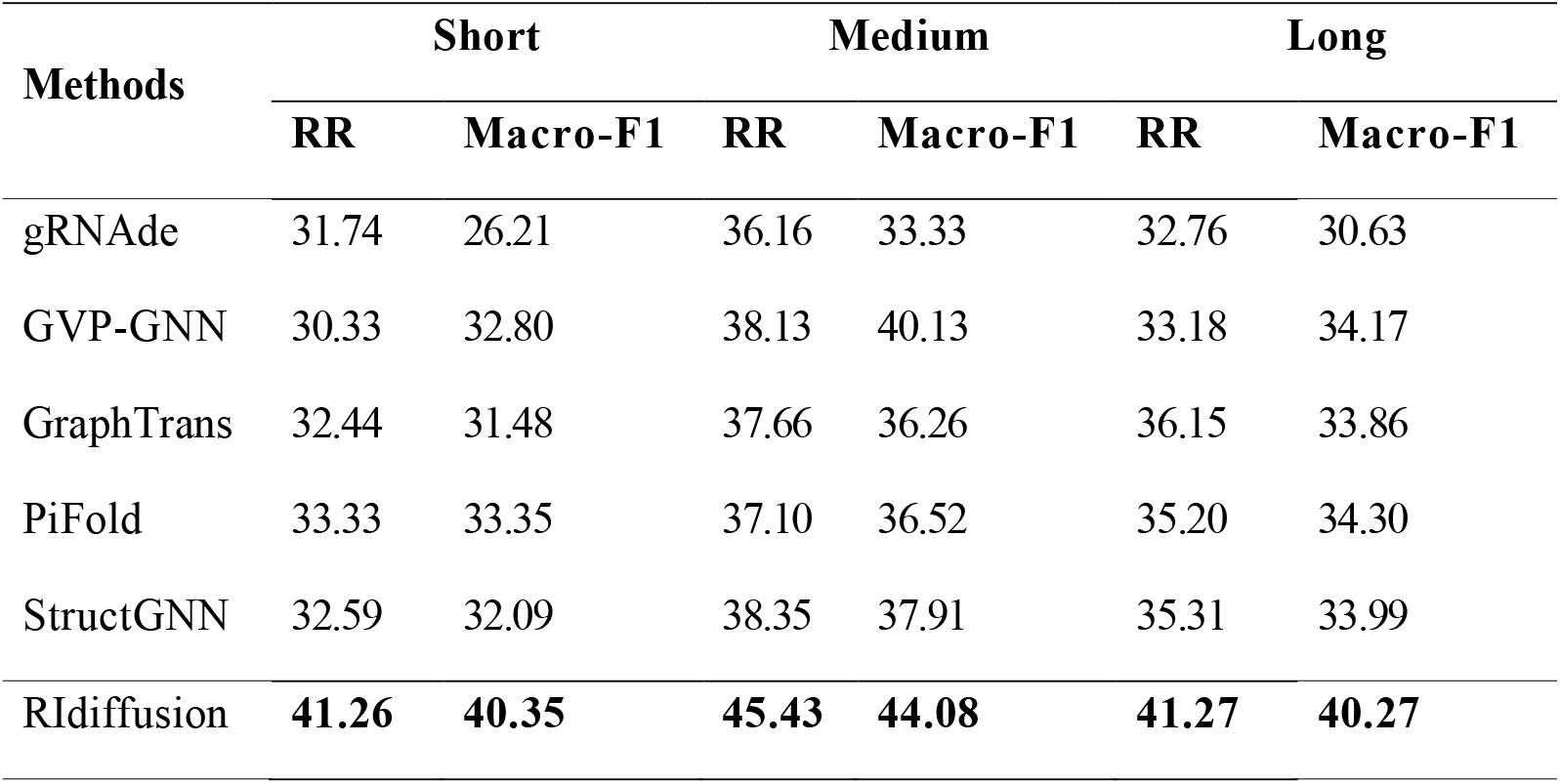
Comparison of recovery rate (RR, %) and Macro-F1 score (Macro-F1, × 100) at a sequence similarity level of 0.8 for sequences of different lengths.

### Novelty and diversity of generated RNA sequences for desired 3D structure

The novelty of generative designs can be viewed from two perspectives: sequence novelty and physicochemical properties novelty.^64,65^ In our case, sequence and physicochemical properties (derived from sequence) novelty are important in RNA 3D inverse folding. Indeed, sequence features such as physicochemical properties play a crucial role in functional RNA design. For instance, it was reported that natural RNAs show privileged physicochemical property space which is related to their biological functions.^66–68^ Other sequence features like GC content are important properties related to developability of RNA molecules. Given the nature of one-to-many relationship between the targeted RNA structure and its possible sequences, the novelty and diversity of generated sequences are important considerations for improving physicochemical properties of the designed sequences.

To assess sequence novelty, we used global-scale novelty (GSN) score, which is the mean novelty scores of structure-guided samples calculated using sequences with maximum, median, and minimum RRs respectively (see Evaluation Metrics in Methods). The GSN indicates the novelty of designs in the whole training sequence space, with higher GSN values reflecting greater novelty. RIdiffusion exhibited the highest novelty score across all RR levels (Table 3). Notably, at the minimum recovery level, our model outperformed other algorithms by a significant margin. This result demonstrates that our model can explore a broader sequence space with promising novelty.

**Table 3.**
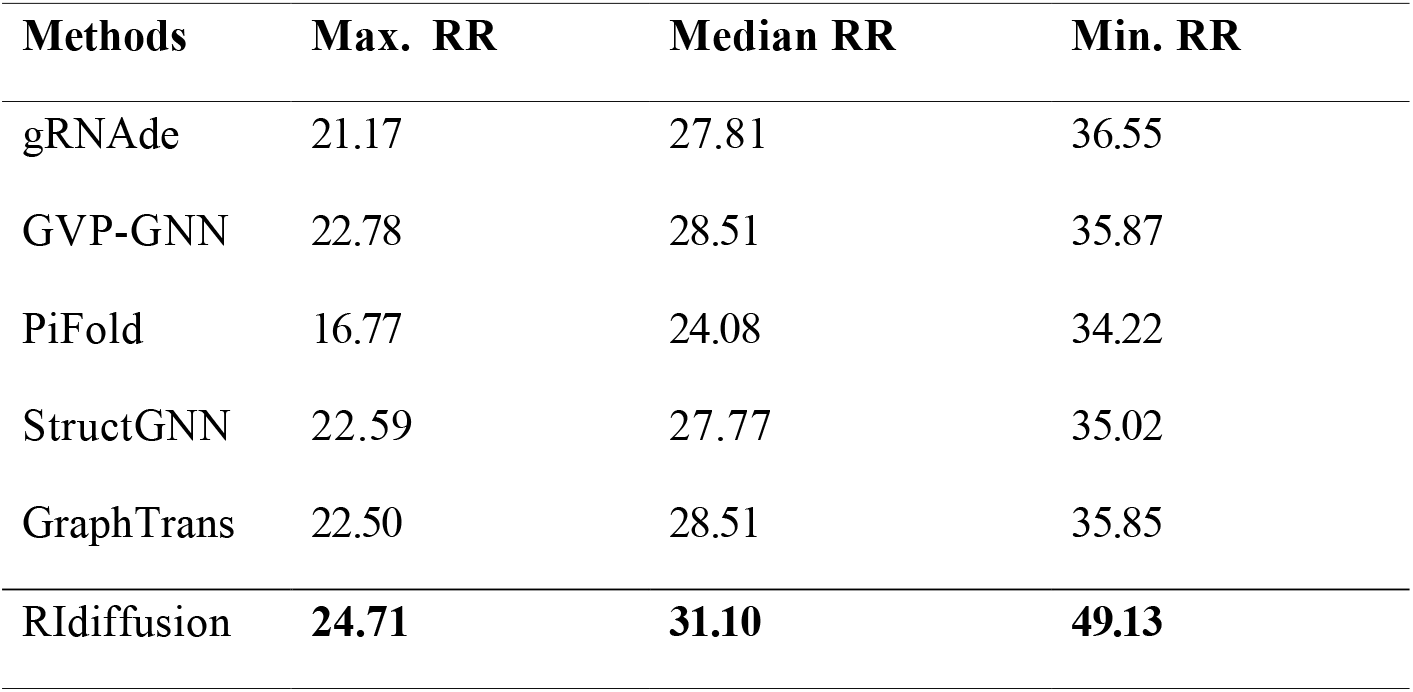
GSN score of generated sequences by different models trained by seq-0.8 dataset.

In addition, we investigated the novelty of physicochemical properties by comparing diverse physicochemical properties (calculated by RNAincoder^56^) for new designs and original sequences using t-SNE dimension reduction visualizations. To assess novelty at varying levels of RR, we classified the sequences into three categories based on recovery rate distribution across various algorithms: low recovery (0∼30%), medium recovery (30∼45%), and high recovery (45∼100%). A general hypothesis is that generated sequences with relatively-high RR have higher chance to be functionally-like to original sequences, and thus their distribution should be different from those generated sequences with lower RR. In another words, if the generative model works meaningfully, the distribution of generated sequences from low recovery, medium recovery and high recovery subsets in the physicochemical properties space should intend to be separated, but not converged. Notably, the samples generated by our model covered a wide range of the physicochemical property space, particularly for sequences with medium and high RRs (Figure 3). This highlights our model’s ability to explore a broader and more novel physicochemical space. In contrast, several baseline models (GraphTrans, GVP, StructGNN, gRNAde) primarily learned limited patterns confined to a few clusters in the t-SNE map, and PiFold failed to exhibit clear clustering trends due to the lack of generated sequences with medium and high RRs.

**Figure 3.**
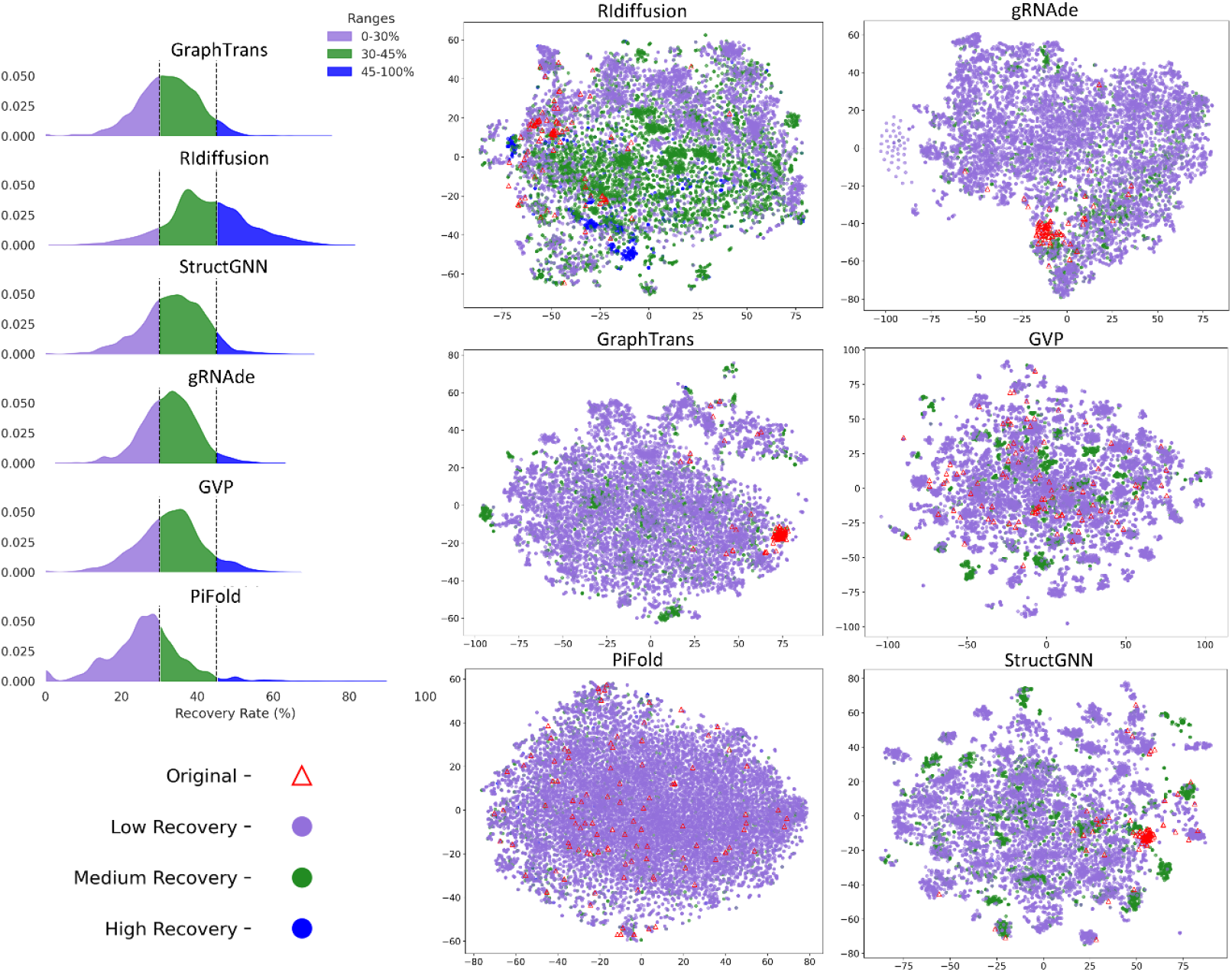
Physicochemical novelty of sequences. Distribution of designed RNA sequences in the physicochemical properties space (mapped using t-SNE).

### Foldability of RNA Sequences Generated by RIdiffusion for Desired 3D Structures

To verify whether the sequences generated by our model can truly fold into the given structure, we used the 3D structure prediction model Rhofold^19^ to predict the structures of the generated sequences. In addition to the quality of sequence redesign, the folding method itself may also introduce potential errors. Therefore, we applied the same structure prediction model to predict the structures of natural sequences and then compared the resulting structures with those of the generated sequences in several case studies (Figure 4). As shown in Figure 4A, RIdiffusion demonstrated the best performance for several structures (5ZEJ, 4V4Z, and 4WR6), exhibiting the lowest RMSD distributions. Furthermore, in three cases (8PIB, 4V4Z, and 4WR6), RIdiffusion generated samples with an RMSD below 2 Å. Meanwhile, typical RNA sequences redesigned by RIdiffusion folded into conformations closely resembling the original input structure (Figure 4B), demonstrating the foldability of sequences generated by our model.

**Figure 4.**
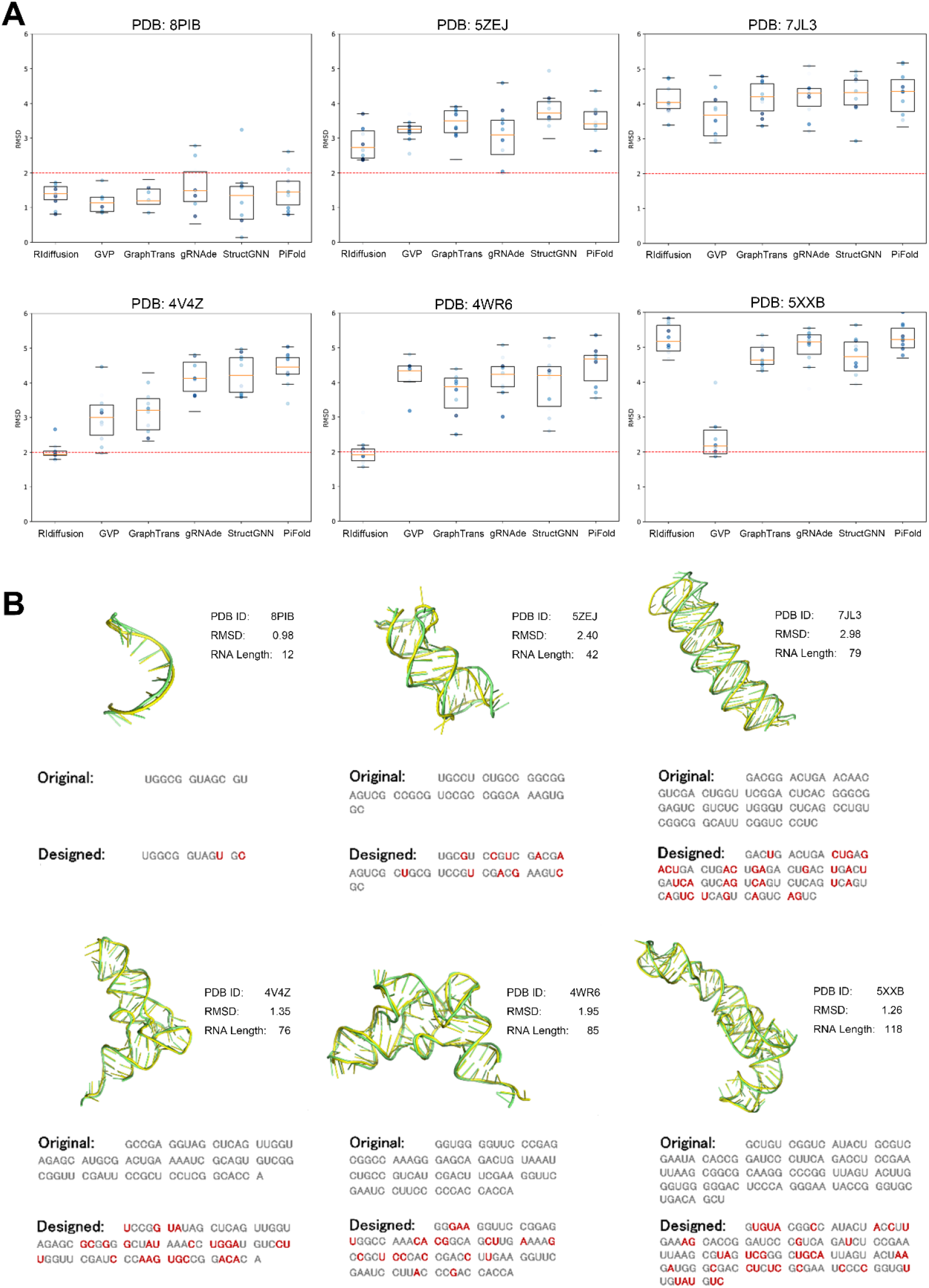
Foldability evaluation of RIdiffusion and other models using Rhofold.^19^ (A) Boxplots showing RMSD distribution of designed sequences generated by inverse folding models. (B) Representative 3D structures of RNA sequences designed by RIdiffusion. The RMSD between the predicted 3D structure of the generated sequence (green) and the original sequence (yellow) was calculated. The red nucleotide indicates sites that differ from the original sequences.

### The contribution of secondary structure (SS) information to 3D design performance is context-dependent

The SS of RNA contains rich biological information and is crucial for accurately predicting the function, stability, and interactions of RNA molecules. It also serves as an important constraint in sequence generation. By integrating SS information, models may better capture key structural features, thereby enhancing prediction accuracy. However, due to the dynamic nature of RNA tertiary structures, static structures derived from PDB may not fully capture conformational variations. To enrich structural information, we incorporated SS predictions from multiple algorithms. We applied three representative RNA SS prediction methods to infer nucleotide contact information, annotated the types of secondary structures, and incorporated the SS data as one-hot encoded node features. Our results indicate that SS information improved RR and Macro-F1 scores of RIdiffusion on the medium-length sequence subset of seq-0.8 dataset (Table 4). However, similar improvements were not observed on short and long subsets of seq-0.8 dataset.

**Table 4.** Performance of RIdiffusion with/without SS information integration on medium-length subsets.

| **Models w/wo SS** | **RR** | **Macro-F1** |
| --- | --- | --- |
| Model 1: without SS | 45.43 | 44.08 |
| Model 2: with SS (RNAstructure) | 46.16 | 44.68 |
| Model 3: with SS (EternaFold) | 45.91 | 45.24 |
| Model 4: with SS (MultiRNAFold) | 46.55 | 45.87 |

### Ablation study of model components highlighting the importance of hyperbolic embedding and HEGNN transformation

To validate the effectiveness of the methods in RIdiffusion, we performed three ablation experiments: 1) We replaced the hyperbolic space representation with Euclidean space representation and substituted HEGNN with EGNN. 2) We used raw 3D positional coordinates as the representation of positional data. 3) We simultaneously applied the replacements described in 1) and 2) to RIdiffusion. Notably, models with hyperbolic embeddings showed superior performance in terms of RR and Macro-F1 score, even with fewer embedding dimensions (see Table S5). Furthermore, results showed that the Transformer layer effectively captured the positional relationships, and hyperbolic embedding significantly enhanced the information representation capabilities of nodes in the graph, collectively improving the performance of our model. (see Table 5).

**Table 5.**
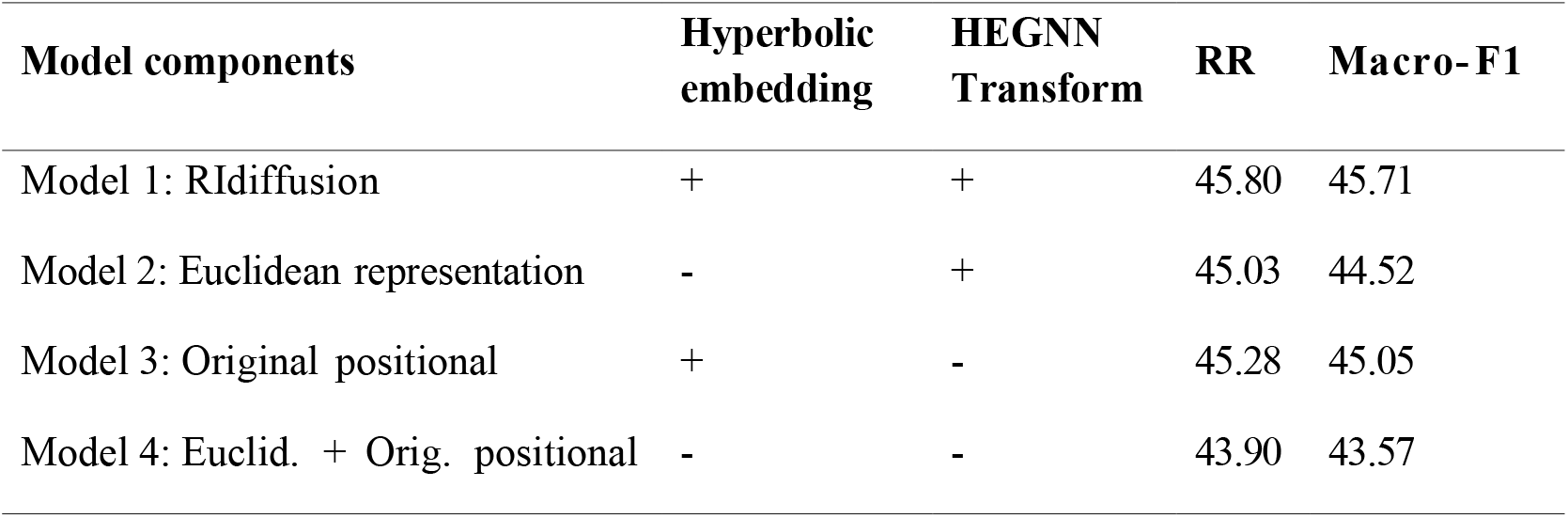
The results of the ablation study of our model, recovery rate (RR, %) and Macro-F1 score (Macro-F1, × 100)

## DISCUSSION AND CONCLUSION

In this study, we present RIdiffusion, a geometric generative model for RNA 3D structure inverse folding. By employing methods such as hyperbolic embedding and HEGNN, we effectively captured the complex structural and geometric properties of RNA molecules, enabling the recovery of the correct sequence from noise. Our results demonstrated that using a discrete diffusion model with geometric deep learning methods can significantly improve the RR of RNA inverse folding and the accuracy of structural design, outperforming baseline methods in terms of RR. Therefore, we hypothesize that embedding RNA structural information in a hyperbolic space may improve the learning efficiency of diffusion models on RNA 3D data.

To comprehensively evaluate our model, we created multiple datasets by partitioning the PDB and RNAsolo datasets based on sequence length and sequence similarity thresholds, following RDesign’s splitting methods while applying even stricter criteria. Experimental results showed that even on low-sequence-similarity datasets (seq-0.8 and seq-0.9) with limited numbers of training samples, our model still outperformed the second-best model by 7.27% and consistently surpassed both benchmark and SOTA models across multiple benchmarks.

In this paper, our main contributions are as follows: We proposed a novel hyperbolic denoising network (HEGNN) to capture the hierarchical structure of RNA and enhance structural similarity measurement for 3D inverse folding, particularly in data-scarce scenarios. We used different computational tools to predict secondary structures as a model enhancement strategy, tested the model and conducted experimental validations. Extensive experiments on datasets with different partitions were conducted to evaluate the generalization capabilities. The results show that our method surpasses existing approaches in both RR and Macro-F1 scores while also generating more novel samples, highlighting the effectiveness of our approach. In the future, we will continue applying this method to various functional RNA design scenarios, incorporating data augmentation techniques such as utilizing *in silico* predicted RNA tertiary structures and embedding relevant biological functional information from high-throughput experiments to enhance downstream tasks in RNA design.

## ASSOCIATED CONTENT

### Data Availability Statement

The original RNAsolo dataset can be accessed via https://rnasolo.cs.put.poznan.pl. The code for discrete diffusion model and hyperbolic network used in this manuscript were adapted from GitHub repositories: https://github.com/cloneofsimo/d3pm.git, https://github.com/HazyResearch/hgcn.git, and https://github.com/ykiiiiii/GraDe_IF.git.

The source codes of RIdiffusion can be accessed at GitHub repository https://github.com/byternaAdmin/RIdiffusion. All datasets and split subsets used in this study were deposited at https://doi.org/10.6084/m9.figshare.28561958.v2.

### Supporting Information

The Supporting Information is available free of charge at https://doi.org/10.6084/m9.figshare.28561958.v2. This includes detailed information on the graph representation of folded RNA, descriptions of datasets, additional experiments comparing Euclidean and Hyperbolic embeddings across various dimensions, and implementation details.

## Author Contributions

X.Z., X.W., D.J., and X.H. provided continuous supervision throughout the project. D.H., S.Z. and M.M. collaboratively conducted the experiments and analysis. S.Z., D.H., Z.W., and X.Z. contributed to the manuscript writing. H.L., R.Z., and H.Z. contributed to data processing and the review of the manuscript. All authors actively participated in result discussions and contributed to the finalization of the manuscript. All authors have given approval to the final version of the manuscript. # These authors contributed equally.

## Funding Sources

This work was supported by the Byterna Therapeutics.

## Notes

Hui Zhao and Shuai Zhang have filed a patent (CN119339781) on key algorithm of this work, the remaining authors declare no competing financial interest.

## ACKNOWLEDGEMENTS

We thank Prof. Xin Guo, Dr. Xuyang Liu, and Dr. Qi Si from Shanghai Academy of AI for Science (SAIS), for their insightful discussions.

## ABBREVIATIONS

RR: Recovery Rate
SOTA: State-of-the-Art
HEGNN: Hyperbolic Equivariant Graph Neural Network
PDB: Protein Data Bank
RNA: Ribonucleic Acid
t-SNE: t-distributed Stochastic Neighbor Embedding
GSN: Global-Scale Novelty
SASA: Solvent Accessible Surface Area
RBF: Radial Basis Function
MHA: Multi-Head Attention
FFN: Feedforward Neural Network
EGNN: Equivariant Graph Neural Network
SE(3): Special Euclidean Group in 3D
SO(3): Special Orthogonal Group in 3D
E(3): Euclidean Group in 3D
DDIM: Denoising Diffusion Implicit Model
CD-HIT: Cluster Database at High Identity with Tolerance
RMSD: Root-Mean-Square Deviation.

## Notes

https://github.com/byternaAdmin/RIdiffusion

https://doi.org/10.6084/m9.figshare.28561958.v2

### Competing Interest Statement

The authors have declared no competing interest.

