## Supplementary Information for "A Hyperbolic Discrete Diffusion 3D RNA Inverse Folding Model for functional RNA design"

### Graph Representation of Folded RNA

RNA is represented as composed of nucleotides, and we represent RNA as a $k$-NN ($k$=10) graph derived from nucleotides, where each residue is represented by three atoms: P, C4', and N1 (pyrimidine) or N9 (purine). We can also construct local coordinates $\mathbf{Q}_i = [\mathbf{u}_i, \mathbf{n}_i, \mathbf{v}_i]$ for each residue, where $\mathbf{u}_i$ is unit vector pointing from C4' atom to P atom, $\mathbf{b}_i$ is unit vector pointing from C4' atom to N1/N9 atom, the normal of this plane $\mathbf{n}_i = \frac{\mathbf{u}_i \times \mathbf{b}_i}{\|\mathbf{u}_i \times \mathbf{b}_i\|}$, and $\mathbf{v}_i = \mathbf{n}_i \times \mathbf{u}_i$. Based on this, we can build translation-invariant features(see Figure S1 and Table S1), which include:

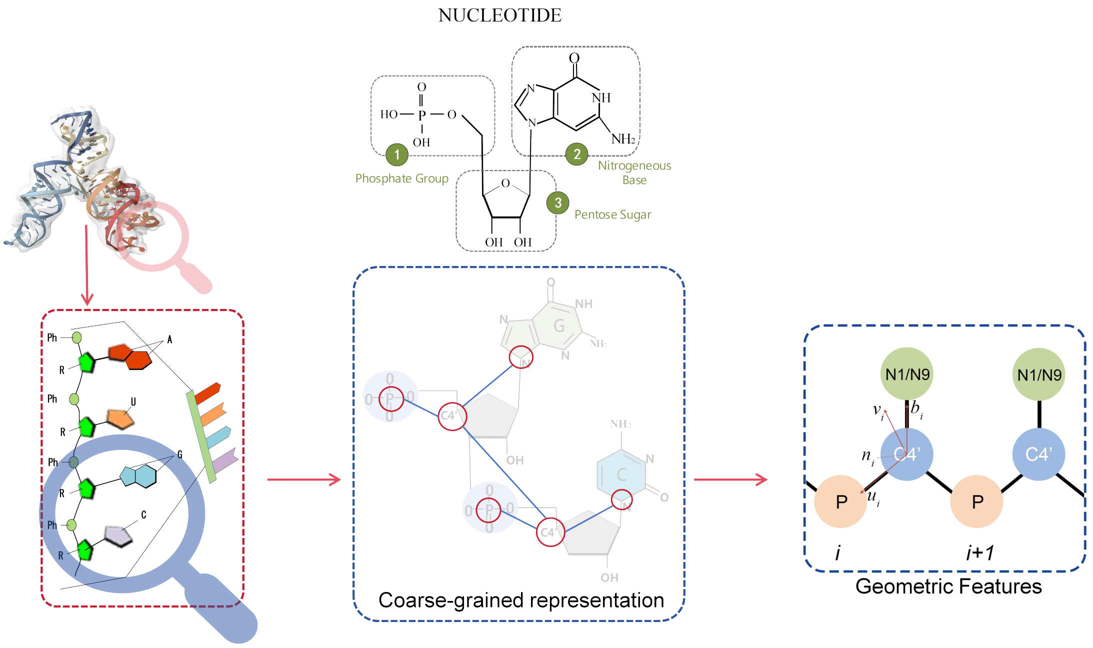


Figure S1: RNA backbone atoms and local coordinates system.

1. **Dihedral Angle:** Given the $i$-th and $(i+1)$-th residue, we describe the dihedral angles as follows:

$$\phi_i = \angle(N_i - P_i - C4'_i - N_{i+1}) \tag{1}$$

$$\psi_i = \angle(P_i - C4'_i - N_i - P_{i+1}) \tag{2}$$

$$\gamma_i = \angle(C4'_i - N_i - P_i - C4'_{i+1}) \tag{3}$$

2. **Distance-Based Edge Features**: For a given pair of atoms, their distance features are represented by a radial basis function (RBF):

$$\text{RBF}(\mathbf{x}_i^{\text{pos}}, \mathbf{x}_j^{\text{pos}}) = \exp\left(-\frac{\|\mathbf{x}_i^{\text{pos}} - \mathbf{x}_j^{\text{pos}}\|^2}{2\sigma^2}\right). \tag{4}$$

   where $\sigma$ represents the standard deviation of the Gaussian function, controlling the width of the bell curve and thus determining the influence range of the distance measure.

3. **Relative Position Edge Features**: These describe the distance between atom $i$ and atom $j$.

4. **Relative Orientation Edge Features**: Each residue has a direction and carries important collective information, represented by the relative orientation $\mathbf{Q}$ in the local coordinate system.

5. **Binary Contact Signal**: This determines whether two residues are in contact in space. If they are in contact, the signal is 1; otherwise, it is 0.

6. **Surface Aware Node Features**: We use the five different scales of surface-aware node features $\mathcal{S}(\mathbf{x})$ defined in Equation 5 and Equation 6 to differentiate the positions of residues in space.

$$\mathcal{S}(\mathbf{x}_i) = \frac{\|\sum_{j \in \mathcal{V}(i)} w_{ij}(C4'_i - C4'_j)\|}{\sum_{j \in \mathcal{V}(i)} w_{ij}^{(k)} \|C4'_i - C4'_j\|}, \tag{5}$$

   where the weights are defined as follows:

$$w_{i,i',\sigma} = \frac{\exp\left(-\frac{\|C4'_i - C4'_j\|^2}{\sigma}\right)}{\sum_{i' \in \mathcal{V}_i} \exp\left(-\frac{\|C4'_i - C4'_j\|^2}{\sigma}\right)} \tag{6}$$

with $\sigma \in \{1, 2, 5, 10, 30\}$.

Furthermore, we also calculated (solvent-accessible surface area)SASA and B-factor to describe the physicochemical properties of RNA molecules.

Table S1: The feature construction of RNA graph structure.

| **Feature** | **Description** |
|---|---|
| Dihedral Angle | $\{\sin, \cos\} \times \{\phi_i, \psi_i, \gamma_i\}$ |
| Distance | $\text{RBF}(\|\text{P}_i - \text{C4}'_i\|)$, $\text{RBF}(\|\text{N1/N9}_i - \text{C4}'_i\|)$ |
| Relative Orientation | $\mathbf{q}(\mathbf{Q}_i^T \mathbf{Q}_j)$ |
| Relative Position | $\mathbf{Q}_i \|\mathbf{C4}'_j - \mathbf{C4}'_i\|$ |
| Contact signal | $\|\text{C4}'_i - \text{C4}'_j\| < 8\text{Å}$ |
| Surface Aware | $\mathcal{S}(\mathbf{x}_i)$ |

### Descriptions of Datasets

#### RNAsolo datasets

RNAsolo is an automatically updating database specifically designed for RNA bioinformatics. It systematically collects experimentally determined RNA three-dimensional structures stored in the Protein Data Bank (PDB), cleans them from non-RNA chains, and groups them into equivalence classes (details are shown in S2 and S3). The database currently includes three types of RNA: standalone RNA molecules, RNA from protein-RNA complexes, and RNA from DNA-RNA hybrids. RNAsolo provides a web service that facilitates user access and data exchange. We followed the splitting methods with structures derived from PDB and RNAsolo described in RDesign,[1] and divided the dataset (2217 structures) into training (1773 structures), validation (221 structures), and test sets (223 structures) in a ratio of 8:1:1.

Table S2: The number of 3D structures of representatives in the RNAsolo database.

| Resolution | Solo RNA | protein–RNA | DNA–RNA |
|---|---|---|---|
| $\leq 1.5$ | 62 | 53 | 7 |
| $\leq 2.0$ | 141 | 267 | 14 |
| $\leq 2.5$ | 203 | 701 | 15 |
| $\leq 3.0$ | 295 | 1380 | 21 |
| $\leq 3.5$ | 321 | 2056 | 21 |
| $\leq 4.0$ | 327 | 2364 | 21 |
| $\leq 20.0$ | 377 | 2767 | 21 |
| all | 567 | 2833 | 25 |

Table S3: The number of 3D structures of all members in the RNAsolo database.

| Resolution | Solo RNA | protein–RNA | DNA–RNA |
|---|---|---|---|
| $\leq 1.5$ | 283 | 491 | 27 |
| $\leq 2.0$ | 647 | 4441 | 39 |
| $\leq 2.5$ | 939 | 8645 | 43 |
| $\leq 3.0$ | 1239 | 11079 | 69 |
| $\leq 3.5$ | 1284 | 12661 | 69 |
| $\leq 4.0$ | 1296 | 13214 | 69 |
| $\leq 20.0$ | 1368 | 13820 | 69 |
| all | 1584 | 13890 | 74 |

### Additional Experiments

#### Implementation Details

We used PyTorch-Geometric (version 2.6.1) and PyTorch (version 2.5.0) for programming and executed our computations on an NVIDIA Tesla V100 GPU.

#### The contribution of secondary structure (SS) information to 3D design performance is context-dependent

Figure S2 and Table S4 compares the results with and without secondary structure (SS) information. Our findings show that incorporating SS information enhanced both the recovery rate (RR) and Macro-F1 scores of RIdiffusion on the medium sequence subset of the seq-0.8 dataset. However, this improvement was not observed in the short and long subsets of the seq-0.8 dataset.

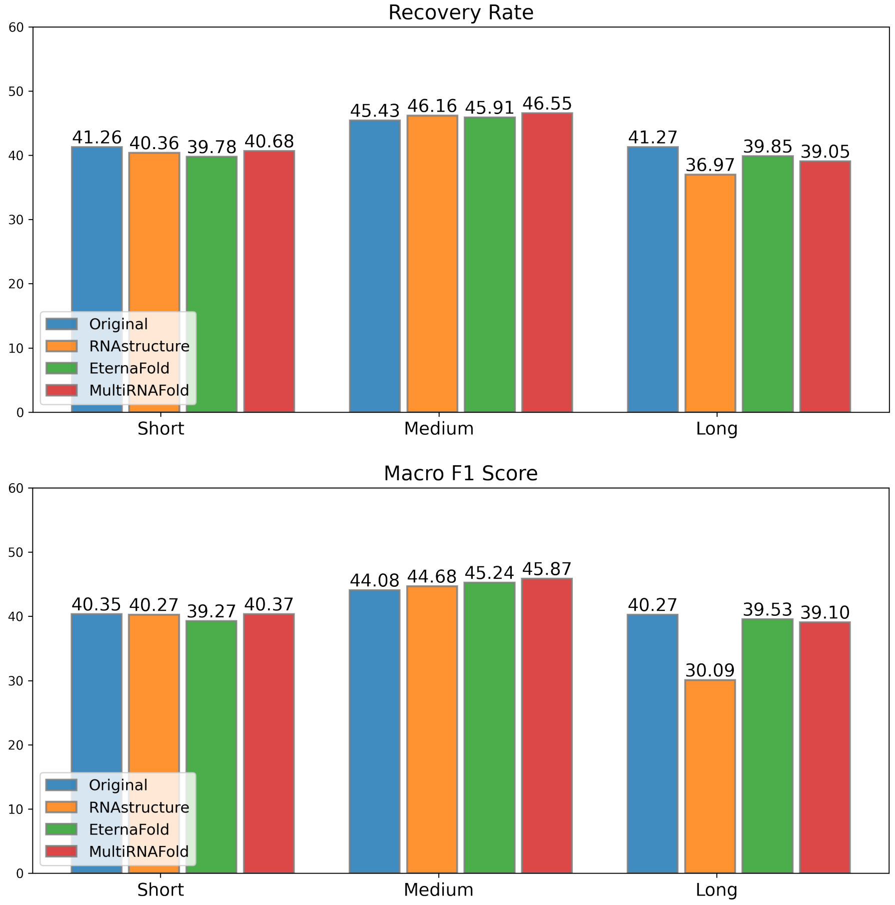


Figure S2: Comparison of different secondary structure prediction enhancement methods with non-secondary structure methods in terms of recovery rate and Macro F1 score across short medium and long subsets.

Table S4: Performance of RIdiffusion with/without SS information integration on short, medium, and long subsets.

| **Models w/wo SS** | **Short** | | **Medium** | | **Long** | |
|---|---|---|---|---|---|---|
| | **RR** | **Macro-F1** | **RR** | **Macro-F1** | **RR** | **Macro-F1** |
| Model 1: without SS | 41.26 | 40.35 | 45.43 | 44.08 | 41.27 | 40.27 |
| Model 2: with SS (RNAstructure) | 40.36 | 40.27 | 46.16 | 44.68 | 36.97 | 30.09 |
| Model 3: with SS (EternaFold) | 39.78 | 39.27 | 45.91 | 45.24 | 39.85 | 39.53 |
| Model 4: with SS (MultiRNAFold) | 40.68 | 40.37 | 46.55 | 45.87 | 39.05 | 39.10 |

#### Additional Ablation Study

In hyperbolic space, the circumference and area of a circle grow exponentially rather than geometrically.[2] Therefore, the distance metric in hyperbolic space increases faster than in Euclidean space[3](see Figure S3), allowing RNA molecules to more compactly represent the geometric relationships of interactions (such as base pairing) in low-dimensional hyperbolic space. Hyperbolic space can efficiently represent high-dimensional structures in low-dimensional space, thereby reducing distortion during the embedding process. This means that we can use lower-dimensional embedding vectors to represent RNA molecules, thereby reducing computational complexity and storage requirements while preserving important structural information. To demonstrate the effectiveness of hyperbolic embeddings, we removed the transformer layers and reduced the number of EGNN layers to weaken the model's representation capability. we conducted comparative experiments of hyperbolic and Euclidean embeddings in different low-dimensional spaces, with the results reported in Table S5.

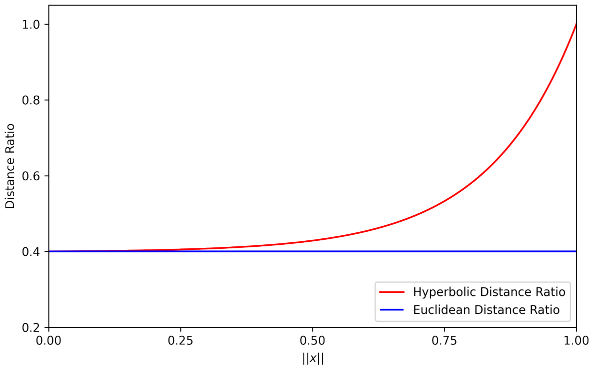


Figure S3: Distances ratio in the Hyperbolic space. Given 3 points,the origin $O$, and points $x$ and $y$ ($\|x\| = \|y\|$). The ratio $\frac{d_E(x,y)}{d_E(x,0)+d_E(0,y)}$ is constant with respect to the movement of points $x$ and $y$ in flat Euclidean space. In contrast, the ratio $\frac{d_H(x,y)}{d_H(x,0)+d_H(0,y)}$ approaches 1 in hyperbolic space, illustrating that the rate of distance growth in hyperbolic space far exceeds that in Euclidean space.

Table S5 shows the comparison of the recovery rates and RIdiffusion F1 scores in the Euclidean space and the hyperbolic space. In low-dimensional spaces, the recovery rate of hyperbolic embeddings consistently outperformed that of Euclidean embeddings, and the 16-dimensional hyperbolic space embedding outperformed the 64-dimensional Euclidean space embedding, indicating that hyperbolic embeddings enhance the model's representation capability.

Table S5: Comparison of Euclidean(euc) Embedding and Hyperbolic(hyp) Embedding.

| **Dimension** | 4 | 8 | 16 | 32 | 64 |
|---|---|---|---|---|---|
| $\text{Recovery}_{hyp}$ | **35.94** | **36.04** | **40.20** | **40.87** | **41.00** |
| $\text{Recovery}_{euc}$ | 31.13 | 31.81 | 32.68 | 32.70 | 36.36 |
| $\text{Macro F1}_{hyp}$ | **36.08** | **36.16** | **39.41** | **39.14** | **40.54** |
| $\text{Macro F1}_{euc}$ | 18.55 | 21.29 | 31.85 | 27.81 | 35.92 |
